# Dissecting the cell cycle regulation, DNA damage sensitivity and lifespan effects of caffeine in fission yeast

**DOI:** 10.1101/2022.11.08.515652

**Authors:** John-Patrick Alao, Juhi Kumar, Despina Stamataki, Charalampos Rallis

## Abstract

Caffeine is a widely consumed neuroactive substance. It can modulate cell cycle progression, override DNA damage checkpoint signalling and increase chronological lifespan (CLS) in various model systems. Early studies suggested that caffeine inhibits the phosphatidylinositol 3-kinase-related kinase (PIKK) Rad3 to override DNA damage-induced cell cycle arrest in fission yeast. We have previously suggested that caffeine modulates cell cycle progression and lifespan by inhibiting the Target of Rapamycin Complex 1 (TORC1). Nevertheless, whether this inhibition is direct or not, has remained elusive. TORC1 controls metabolism and mitosis timing by integrating nutrients and environmental stress response (ESR) signalling. Nutritional or other stresses activate the Sty1-Ssp1-Ssp2 (AMP-activated protein kinase complex, AMPK) pathway, which inhibits TORC1 and accelerates mitosis through Sck2 inhibition. Additionally, activation of the ESR pathway can extend lifespan in fission yeast. Here, we demonstrate that caffeine activates Ssp1, Ssp2 and the AMPKβ regulatory subunit Amk2 to advance mitosis. Ssp2 is phosphorylated in an Ssp1-dependent manner following exposure to caffeine. Furthermore, Ssp1 and Amk2, are required for resistance to caffeine under conditions of prolonged genotoxic stress. The effects of caffeine on DNA damage sensitivity are uncoupled from mitosis in AMPK pathway mutants. We propose that caffeine interacts synergistically with other genotoxic agents to increase DNA damage sensitivity. Our findings suggest that caffeine accelerates mitotic division and is beneficial for CLS through AMPK. Direct pharmacological targeting of AMPK may serve towards health span and lifespan benefits beyond yeasts, given the highly conserved nature of this key regulatory cellular energy sensor.

## Introduction

The widely consumed neuroactive methylxanthine compound caffeine, has been linked to increased chronological lifespan (CLS), protective effects against diseases such as cancer and improved responses to clinical therapies (Alao and Sunnerhagen, 2020; Bode and Dong, 2007; Cross and Gunter, 2018; Löf et al., 2015; Rallis et al., 2013). Deciphering the mechanisms by which caffeine overrides DNA damage checkpoint signalling has been of particular interest (Alao and Sunnerhagen, 2020). While extensively studied over the last three decades, the precise mechanisms whereby caffeine exerts its activity on cell cycle regulation have remained unclear (Alao et al., 2021, 2020, 2014; Alao and Sunnerhagen, 2020; Bentley et al., 1996).

Eukaryotic cells are subject to continuous DNA damage, resulting from the effects of metabolic processes as well as exposure to environmental agents such as ionizing radiation, UV radiation, cigarette smoke or chemotherapeutic agents (Carusillo and Mussolino, 2020; Ciccia and Elledge, 2010; Paviolo et al., 2019). The *S. pombe* phosphatidylinositol 3-kinase-related serine/threonine kinase (PIKK) Rad3 (a homologue of mammalian ataxia telangiectasia mutated (ATM) and ATM related kinase (ATR)), is activated in response to DNA damage or replication stress. Rad3 activation and phosphorylation creates binding sites for adapter proteins that facilitate the activation of downstream kinases Cds1 and Chk1 in the S phase and G2 phases of the cell cycle, respectively. During S phase Cds1 phosphorylates Cdc25 on serine and threonine residues resulting in its activation and sequestration within the cytoplasm following binding of the 14-3-3-related, Rad24 protein. Rad3 and Cds1 also activate the Mik1 kinase during S phase which directly inhibits Cdc2 (Bentley et al., 1996; Christensen et al., 2000). In *S. pombe*, the reciprocal activity of the Cdc25 phosphatase and Wee1 kinase on Cdc2 activity, determines the actual timing of mitosis. Additionally, environmental cues such as nutrient limitation and environmental stress can influence the timing of mitosis through target of Rapamycin complex 1 (TORC1) signalling (Atkin et al., 2014; Davie et al., 2015). The inhibition of TORC1 activity results in the activation of the Greatwall related-kinases Ppk18 and Cek1. Ppk18 and Cek1 in turn activate the endosulphine Igo1 which inhibits PP2A^pab1^ to induce mitosis (Pab1 being the regulatory subunit in *S. pombe*) (Aono et al., 2019; Chica et al., 2016a).

Initial studies suggested that caffeine inhibits Rad3 and its homologues to override DNA damage checkpoint signalling in both yeast and mammalian cells (Moser et al., 2000; Osman and McCready, 1998; Wang et al., 1999). These findings, based on *in vitro* experiments demonstrating that caffeine can inhibit Rad3 and ATM, have been proven controversial. Firstly, caffeine has been shown to override DNA damage checkpoint signalling without blocking ATM and Rad3 signalling (Alao et al., 2014; Cortez, 2003; Kunoh et al., 2008). Furthermore, caffeine can also inhibit other members of the PIKK family (Sarkaria et al., 1999). More recent studies indicate that caffeine inhibits TORC1 signalling in yeast and mammalian cells. Indeed, TORC1 inhibition in *S. pombe* is sufficient to override DNA damage checkpoint signalling (Alao et al., 2020; Davie et al., 2015). It remains unclear, if caffeine directly inhibits TORC1 by acting as a low intensity ATP competitor or indirectly via the *S. pombe* AMP-Activated Protein Kinase (AMPK) Ssp2 (Reinke et al., 2006; Wanke et al., 2008). Caffeine has also been shown to extend CLS in *S. pombe* and induce in combination with Rapamycin, gene expression patterns similar to nitrogen-starved cells or cells in which *tor2* is conditionally deleted (through a temperature sensitive (TS) mutation). Mutants lacking *ssp2* display a shortened CLS. It remains unclear if caffeine alters gene expression or extends CLS directly, or through Ssp2 activation (Rallis et al., 2017, 2014).

In the current study, we have identified novel roles for the Ssp1 and Ssp2 kinases in facilitating the indirect inhibition of TORC1 by caffeine. Caffeine failed to override DNA damage checkpoint signalling in *ssp1*Δ and *ssp2*Δ mutants. We demonstrate that caffeine interacts with Ssp1 signalling and induces Ssp2 phosphorylation in an Ssp1-dependent manner. We also identify a specific role for Amk2, the β-regulatory subunit of the AMPK complex, in mediating resistance to caffeine, possibly through chromatin remodelling modulation and DNA damage repair. Downstream of TORC1, caffeine induced mitosis in *ppk18*Δ and *cek1*Δ *ppk18*Δ but not *igo1*Δ mutants, but this activity did not correlate with its effect on DNA damage sensitivity. Caffeine may, thus, enhance DNA damage sensitivity independently of mitosis by interfering with DNA damage repair. TORC1 inhibition with Torin1 also potently sensitized *ppk18*Δ, *cek1*Δ *ppk18*Δ and *igo1*Δ mutants to DNA damage. These observations suggest roles for Ppk18, the related kinase Cek1, an unknown kinase and possibly an unidentified probable role for Igo1 in DNA damage repair and/or checkpoint recovery (Amar-Schwartz et al., 2022; Campos and Clemente-Blanco, 2020; Ramos et al., 2019). Exposure to caffeine also suppressed Sck2 expression. Sck2 functions downstream of TORC1 to regulate the timing of mitosis (Chica et al., 2016b) and plays a role in regulating CLS (Roux et al., 2006). We previously demonstrated that caffeine prolongs CLS in *S. pombe* (Rallis et al., 2013). Nevertheless, whether this is due to direct TORC1 inhibition or through stress response induction has remained elusive. In this study, we demonstrate that the caffeine-mediated CLS extension is Ssp1, Ssp2 and Amk2-dependent. Together these findings provide evidence for the Ssp1-AMKP-TORC1 signalling axis in mediating the effects of caffeine on both cell cycle progression and chronological ageing in *S. pombe*.

## Results

### *S. pombe* Ssp1 and Ssp2 mediate the effect of caffeine on the G2 DNA damage checkpoint

As previously reported, both caffeine (10mM) and Torin1 (5µM) override the cell cycle arrest induced by 5µg/mL phleomycin in wild type *S. pombe* cells. In contrast, cells exposed to phleomycin or left untreated remained arrested while control cultures proliferated with steady state kinetics (Figure 1A) (Alao et al., 2020). We hypothesized that caffeine overrides DNA damage checkpoint signalling at least partially, via inhibition of TORC1 activity (Alao et al., 2020). Direct inhibition of TORC1 activity with Torin1 is more effective than exposure to caffeine raising the possibility that the latter may act indirectly. We, therefore, wondered about the implication of AMPK and investigated the possible role of Ssp1 and Ssp2 in modulating the mitotic effects of caffeine. Nutrient deprivation or exposure to environmental stress indirectly inhibits TORC1 activity, via activation of the Ssp1-Ssp2 (AMPK catalytic (α) subunit) pathway. Deletion of *ssp1* or *ssp2* inhibited the effect of caffeine on DNA damage induced cell cycle arrest in cells exposed to phleomycin (Figure 1B, C). It is likely, that deletion of *ssp1* or *ssp2* affects TORC1 activity and hence the kinetics of Torin1-induced mitosis relative to *wt* cells. In contrast, the effect of Torin1 under similar conditions was partially suppressed (or dampened) but not inhibited until after 120 min. TORC1 inhibition leads to activation of the Greatwall homologue Ppk18 which activates Igo1. Igo1 in turn, inhibits PP2A^Pab1^ activity causing cells to enter mitosis prematurely with a shortened cell size (Aono et al., 2019; Chica et al., 2016b; Davie et al., 2015; Pérez-Hidalgo and Moreno, 2017). Deletion of *ppk18* did not suppress the ability of either caffeine or Torin1 to override DNA damage checkpoints. We noted however, that the *ppk18*Δ mutant reached the maximum level of septation more slowly than wild type cells exposed to phleomycin and caffeine or Torin1 (90 min versus 60 min in *ppk18*Δ and *wt* strains respectively) (Figure 1D). Surprisingly, absence of both *cek1* with *ppk18* (regulating Igo1, (Aono et al., 2019)) suppressed the effects of Torin1 but not caffeine on phleomycin-induced cell cycle arrest (Figure 1E). It should be noted, that Torin1 induces cell cycle arrest while caffeine does not. It is possible that these differential results are due to TORC2 inhibition by Torin1 but not caffeine. In contrast, deletion of *igo1* markedly reduced the ability of both caffeine and Torin1 to override DNA damage checkpoint signalling (Figure 1F).

**Figure 1.**
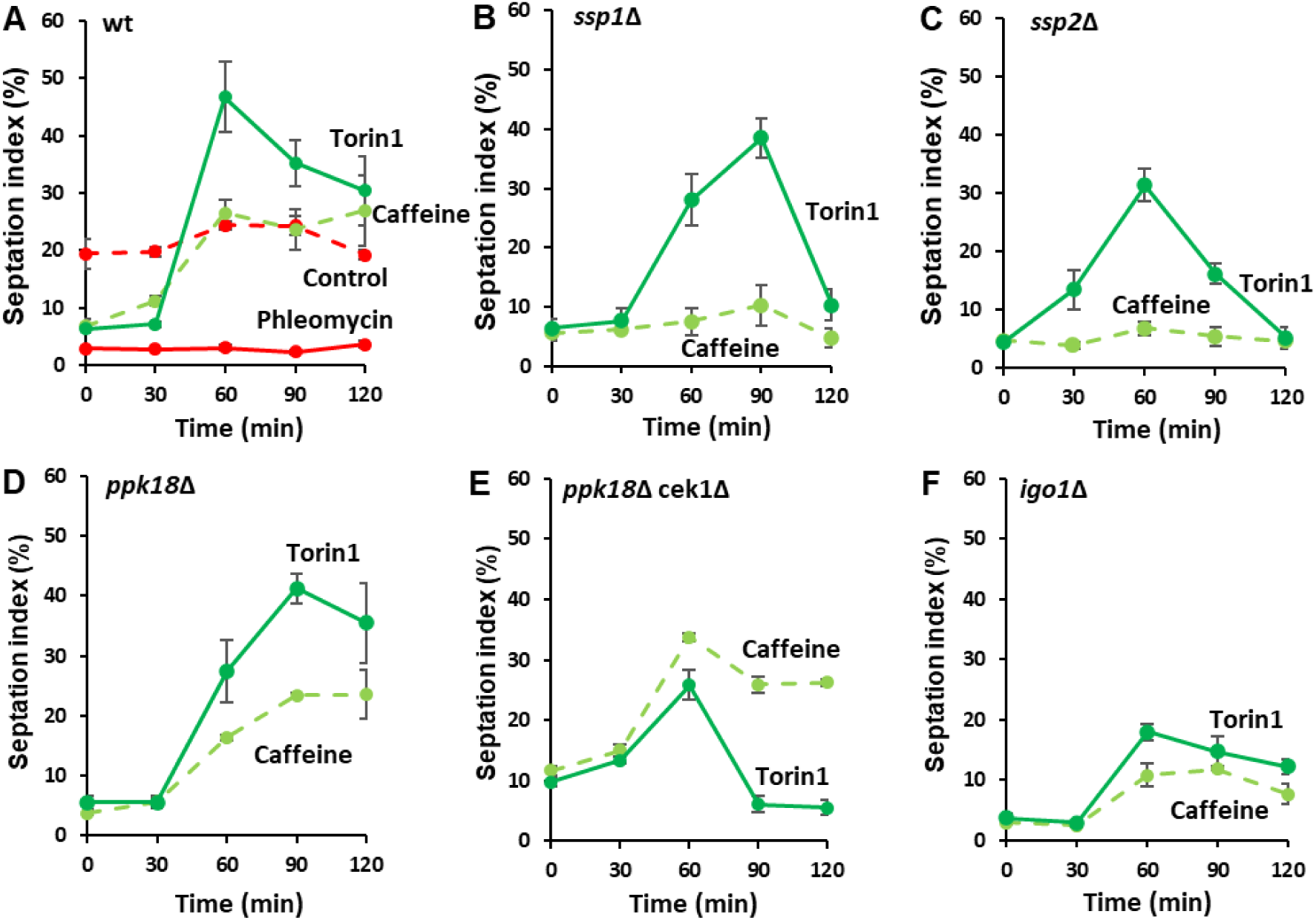
*S. pombe* AMPK mediates the effects of caffeine on cell cycle progression. **(A-F).** Wild type (*wt*) and mutant cells were treated as indicated and harvested at every 30 minutes over a 2h period. Cells were stained with calcofluor and the septation indices determined by fluorescent microscopy

Wild type and *ssp1*Δ strains were sequentially exposed to phleomycin and caffeine or Torin1 and subjected to FACS analysis. No changes in wild type DAPI intensity were observed when cells were exposed to phleomycin alone (1.03% vs 0,83% of cells passing a 40K DAPI intensity arbitrary cutoff in time 0 and 120min respectively and with geometric means of fluorescent intensity 22,755 and 23,329, respectively). An increase in the case of *ssp1*Δ was observed with phleomycin exposure (1.09% vs 6.34% of cells passing a 40K DAPI intensity arbitrary cutoff in time 0 and 120min respectively and with geometric means of 24,581 and 28986 respectively). Exposure of *wt* cells to caffeine in the presence of phleomycin clearly resulted in an increase in DAPI intensity above 40K point (0.69% vs 21.4% of cells above 40K respectively and geometric means 22,611 vs 31,766). This effect was suppressed in *ssp1*Δ mutants where the increase above 40K was from 0.69% to 9.49% (geometric means from 22,897 to 30212). Given that phleomycin alone, induces an increase of DAPI intensity in *ssp1*Δ, the change attributable to caffeine is even lower compared to wild type cells (Supplementary figure 1). No notable changes can be observed for wild type or *ssp1*Δ cells treated with phleomycin and Torin1. Taken together (see also relevant septation index measurements in Figure 1A and 1B), our results may indicate that following phleomycin treatment, caffeine can drive DNA replication and mitosis in an Ssp1-dependent manner. Interestingly, we could not detect differences in DNA damage sensitivity in *wt*, *ssp1*Δ, *ssp2*Δ, *cek1*Δ *ppk18*Δ or *igo1*Δ mutants co-exposed to phleomycin and caffeine or Torin1 (Figure 2, Supplementary Figure 2).

**Figure 2.**
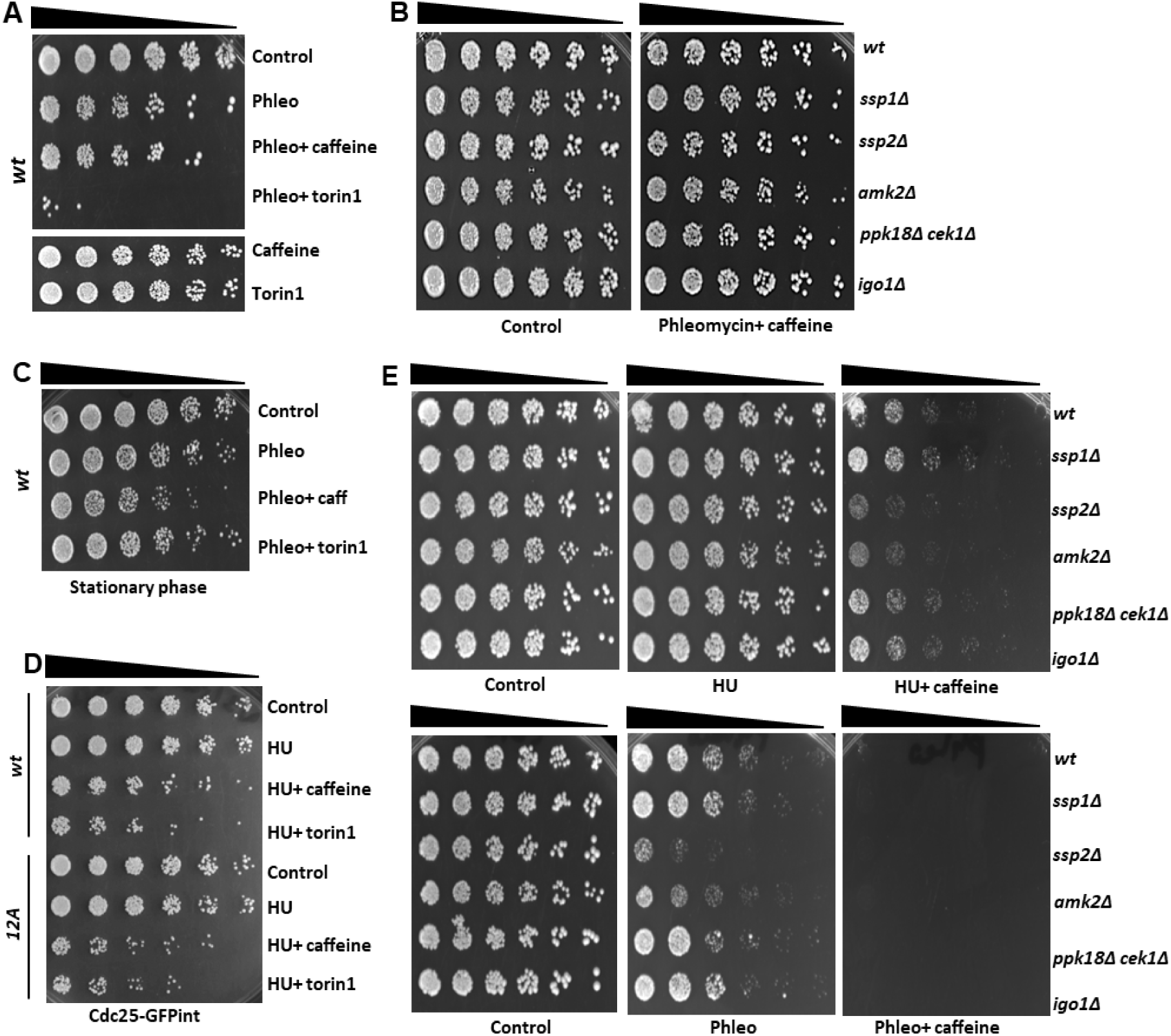
Caffeine enhances DNA damage sensitivity independently of its cell cycle effects. **(A).** The indicated wild type cells were grown to log phase and left untreated or incubated with 5 µg/ mL phleomycin for 2h. Cultures were then incubated for a further 2h with or without 10mM caffeine or 5µM Torin1 in 10mL medium for a further 2h as indicated. Cultures were adjusted for cell number, serially diluted 3-fold, plated on YES agar and incubated for 3-5 days at 32°C. (**B).** Log phase cultures of the indicated strains were treated as in A. **(C).** Wild type cells were grown to stationary phase and exposed to phleomycin for 2h and then cultured with and without caffeine for a further 3 h. Samples were adjusted for cell number and treated as in A. **(D).** Wildtype cells expressing Cdc25-GFP and Cdc25 (12A)-GFP were exposed to 20mM HU for 2h and then incubated with 10mM caffeine for a further 2h and treated as in A. **(E).** Log phase cultures of the indicated strains were plated on YES agar containing 5 mM HU or 5 µM phleomycin with or without 10 mM caffeine.

As previously reported, both caffeine and Torin1 increase sensitivity to phleomycin in liquid culture (Alao et al., 2014) (Figure 2A). We have consistently observed, that Torin1 is more effective at enhancing DNA damage sensitivity. TORC2 is required resistance to DNA damage, hence its inhibition by Torin1 might exert different effects on viability. Although deletion of *ssp1*, *ssp2* and *igo1* abrogated the effect of caffeine on mitosis, we did not detect a reduction in DNA damage sensitivity relative to *wt* cells under these conditions (Figure 1, Figure 2B, Supplementary Figure 2). Both drugs clearly enhanced the sensitivity of *ssp2*Δ, *cek1*Δ *ppk18*Δ, *igo1*Δ mutants to phleomycin, despite the lack of mitotic progression (Figure 1, Figure 2B). While Ssp1 cells may display slightly higher resistance to phleomycin with caffeine, we did not detect a clear relationship between mitosis and viability (Supplementary figure 2A to 2C). Similarly, both caffeine and Torin1 sensitized *igo1*Δ mutants to phleomycin like *wt* cells, despite the marked difference in mitosis (Figure 1F, 2B and supplementary figure 2D and 2E). Caffeine also slightly enhanced the DNA damage sensitivity of *wt* cells exposed to phleomycin in cultures treated in stationary phase (Figure 2C). Thus, caffeine and Torin1 exert differential effects on DNA damage sensitivity in a context dependent manner. As caffeine can increase HU sensitivity in a Cdc25-dependent manner (2D) (Alao et al., 2014), we next examined the effect of caffeine on *wt* and mutant cells under chronic genotoxic conditions. Under these conditions, *ssp2* and *amk2* mutants showed increased sensitivity to HU combined with caffeine. Furthermore, *ssp2* and *amk2* mutants showed increased sensitivity to phleomycin alone while co-exposure to caffeine was lethal to all strains (Figure 2E). Caffeine has been reported to be genotoxic and can enhance DNA damage sensitivity without advancing mitosis. These observations suggest that caffeine may induce a specific form of DNA damage when combined with genotoxic agents (Calvo et al., 2009; Rowley and Zhang, 1999).

### Caffeine induces Ssp1-dependent Ssp2 phosphorylation on Thr169

To further investigate the effect of caffeine on Ssp2 signalling, we used a previously reported phospho-specific antibody (Schutt and Moseley, 2017). Ssp2 phosphorylation was clearly induced in *wt* cells expressing Myc-tagged Ssp2 when exposed to 10mM caffeine for 2h, albeit to a lesser degree than glucose deprivation. Ssp2 phosphorylation was suppressed in cells exposed to 5µM Torin1 under similar conditions (Figure 3A, 3B and 3C). Environmental stresses induce Ssp1-mediated phosphorylation of Ssp2. In contrast to the previously reported transient effect of 1M KCl on Ssp2 phosphorylation (Schutt and Moseley, 2017), this effect was sustained in cells exposed to caffeine over a 2h period (Figure 3C). Caffeine also increased the amount of Ssp2 phosphorylation in cells previously exposed to phleomycin, while co-exposure to Torin1 exhibited the opposite effect (Figure 3D). Although Ssp2 is activated by changes in the ratio of AMP to ATP, it can also be activated by cellular stresses. Exposure to 1M KCl, has previously been shown to suppress cellular ATP levels (Schutt and Moseley, 2017). This suggests that caffeine indirectly inhibits TORC1 via Ssp2 in contrast to Torin1. It is unclear why ATP suppresses sensitivity to phleomycin but supports our contention that viability is independent of mitosis in this context. The effect of caffeine on Ssp2 phosphorylation was largely independent of ATP in cells previously exposed to phleomycin, while Torin1 suppressed phospho-Ssp2 levels. ATP slightly suppressed the caffeine-induced phosphorylation of Ssp2. Additionally, ATP also increased phospho-Ssp2 levels in cells exposed to phleomycin and Torin1 (Figure 3E). Together, these finding suggest that caffeine activates Ssp2 independently of ATP although it may suppress Torin1 activity. ATP did not affect the ability of caffeine or Torin1 to drive phleomycin-arrested cells into mitosis (Figure 3F). We noted however, that septated cells exposed to ATP in the presence of phleomycin and caffeine were longer than cells exposed to the latter alone (Figure 3G). Additionally, ATP suppressed the effect of caffeine and Torin1 on phleomycin sensitivity independently of its effects on cell cycle progression (Figure 3H). Caffeine in contrast to 1M KCl, thus, appears to induce Ssp2 phosphorylation and cell cycle progression independently of ATP.

**Figure 3.**
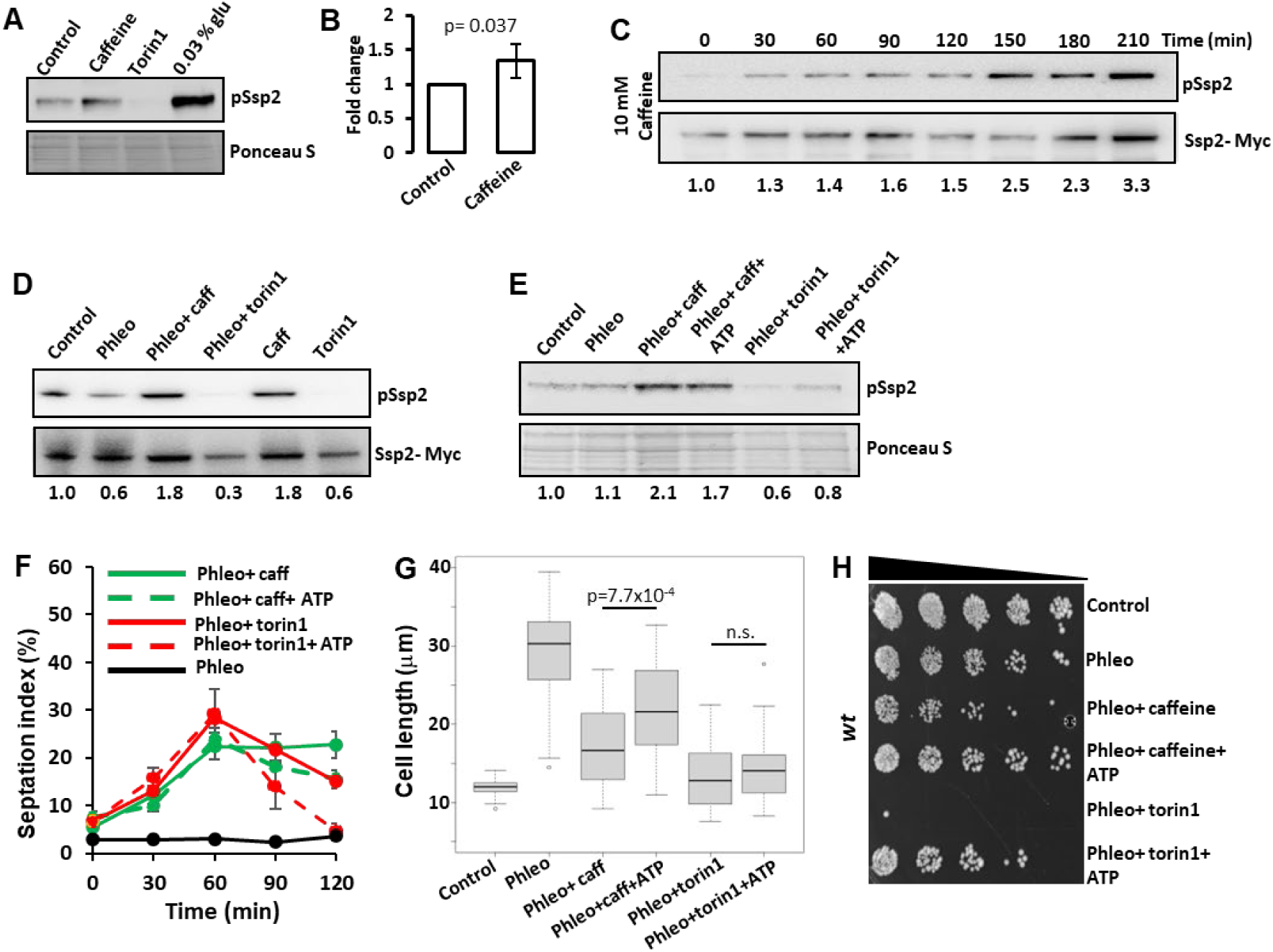
Caffeine induces Ssp2 phosphorylation independently of ATP. **(A).** Log phase wild type cells were exposed to 10 mM caffeine, 5 µM Torin1 or grown in media containing 0.03% glucose for 2 hrs. Samples were resolved by SDS-PAGE and probed with antibodies directed against phospho-Ssp2. Membranes were subsequently stained with ponceau S. **(B).** Blots were analysed using imageJ software. Error bars represent the mean ±S.E. of 3-independent experiments. **(C).** Log phase cells expressing Ssp2-myc, were exposed to 10mM caffeine and harvested at the indicated time points. Samples were resolved by SDS-PAGE and probed with antibodies directed against phospho-Ssp2 and Myc. **(D).** Log phase cells expressing Ssp2-myc, were exposed to 5µg/mL phleomycin for 2 hrs and then exposed further with 10mM caffeine or 5µM Torin1 as indicated. Samples were treated as in A. **(E).** Log phase wild type cells were treated as in D. Cells were pretreated with 10mM ATP for 30 minutes. (**F).** Log phase cells were diluted into fresh media and exposed to 5µg/mL phleomycin for 2 hrs. Cultures were then left untreated or exposed to 10mM caffeine or 5 µM Torin1 with or without 10mM ATP and harvested at the indicated time points. Cells were stained with calcofluor and the septation index determined using fluorescent microscopy. **(G).** Cells were treated as in F and cell length (n= 50) determined. Results were analysed using a student’s t-test with unequal distribution. **(H).** Cells were treated as in F, serially diluted 3-fold and plated on YES agar.

### Caffeine modulates TORC1- regulated mitotic signalling

Indirect inhibition of TORC1 by Ssp2 results in the suppression of Sck2 activity, Ppk18 and Cek1 activation, Igo1 activation and downstream inhibition of PP2A activity leading to mitotic entry (Chica et al., 2016b). We, thus, examined the effect of caffeine on TORC1-regulated cell signalling. Caffeine clearly suppressed Sck2 but not Sck1 (an Sck2 homologue that plays a minor role in regulating the timing of mitosis) expression in a time-dependent manner (Figure 4A and 4B). A similar effect was observed in cells previously exposed to phleomycin, where caffeine may be able to induce the accumulation of Sck1 (Figure 4C) but suppresses Sck2 expression in a time-dependent manner (Figure 4D). Caffeine may thus indirectly induce Cdc25 activation, Wee1 inhibition and consequently, mitosis via suppression of Sck2 expression and, consequently, reduction of its corresponding enzymatic activity (Figure 4B). We did not detect an appreciable level of reduced Maf1 phosphorylation in cells exposed to caffeine alone (Figure 4E). The output of caffeine-mediated TORC1 inhibition, may thus differ from those of Rapamycin and Torin1. Caffeine may affect Cdc25 activity independently of TORC1. We previously reported that caffeine stabilizes Cdc25 in *S. pombe* (Alao and Sunnerhagen, 2020). We note that HU but not phleomycin induces degradation of the Cdc25(12A)-GFPint (Figure 4F and 4G). Inappropriate Cdc25 activity during S-phase may interfere with DNA replication and repair. Caffeine likely drives mitosis via TORC1 inhibition to activate PP2A while simultaneously stabilizing Cdc25 via activation of TORC2 (See below).

**Figure 4.**
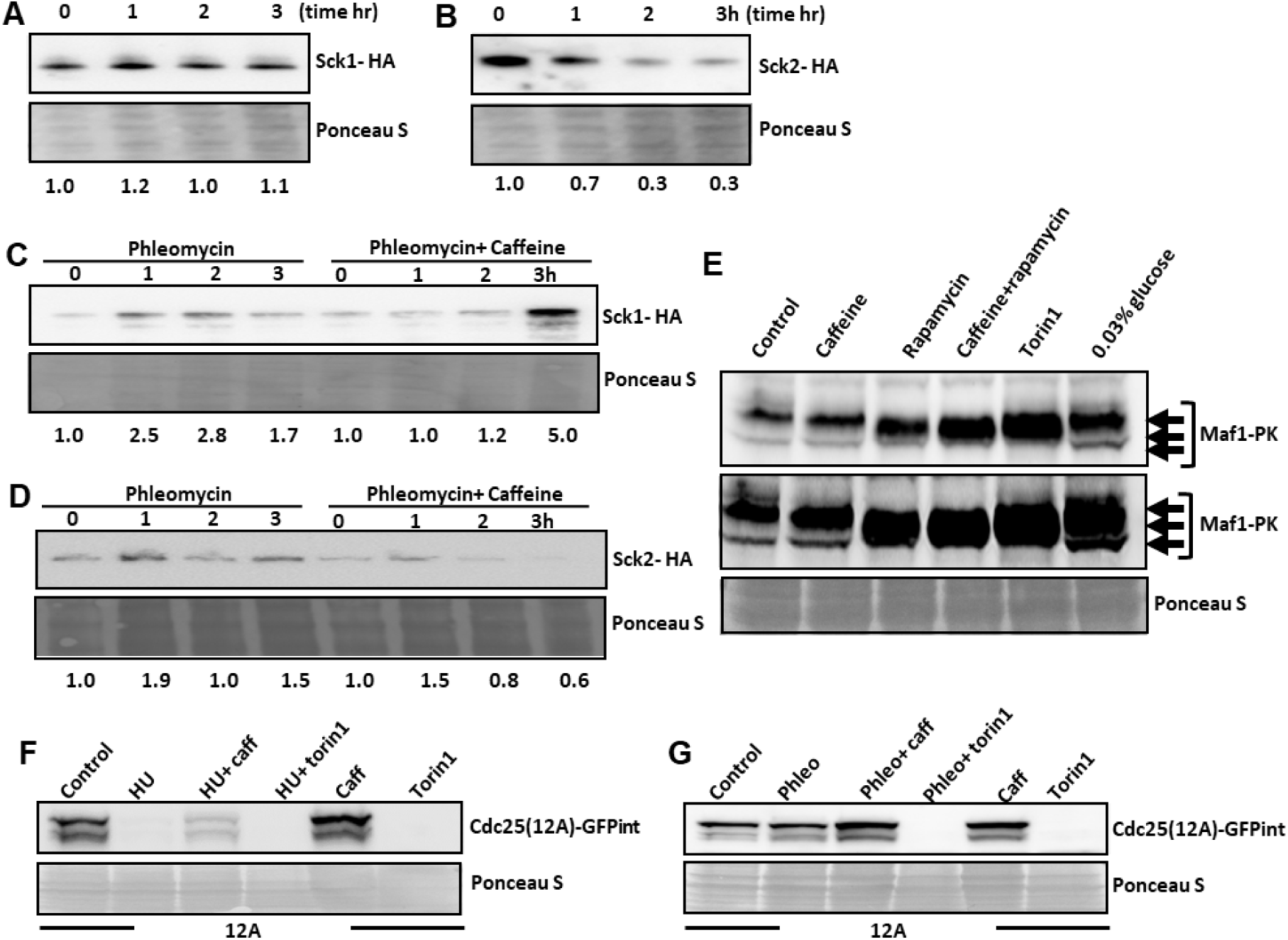
Caffeine indirectly inhibits TORC1 signalling in *S. pombe*. **(A).** Cells expressing Sck1-HA were grown to log phase, resuspended in fresh YES media and exposed to 10mM caffeine. Samples were harvested at the indicated at the indicated timepoints, resolved by SDS-PAGE and membranes probed with anti-HA antibodies. Ponceau S was used to monitor gel loading. **(B).** Cells expressing Sck2-HA were treated as in A. **(C).** Cells expressing Sck1-HA were exposed to 5µg/mL phleomycin for 2h and then further incubated for 2h with or without 10mM caffeine and treated as in A. **(D).** Cells expressing Sck2-HA were treated as in C. **(E).** Cells expressing Maf1-pk were treated with 10 mM caffeine, 100 ng/mL Rapamycin, caffeine and Rapamycin or 5µM Torin1 for 90 minutes and analysed by SDS-PAGE. Membranes were probed with antibodies directed against V5. **(F).** Cells expressing Cdc25-GFP were exposed to 20mM HU for 2h and then for a further 2h with or without 10mM caffeine or 5µM Torin1. **(G).** Cells expressing Cdc25(12A)-GFP were exposed to 5ug/ mL phleomycin for 2h and then for a further 2h with or without 10mM caffeine or 5µM Torin1. Membranes were probed with antibodies directed against GFP.

### Ssp2 mediates the effect of caffeine on cell cycle progression in the absence of checkpoint signalling

We next investigated the role of Ssp2 in mediating the cell cycle effects of caffeine under normal growth conditions. Exposure to 10mM caffeine or 5µM Torin1 advanced *wt* cells into mitosis over a 2 h period (Figure 5A). Deletion of *ssp2* partially suppressed the effect of both caffeine and Torin1 on cell cycle progression (Figure 5B, for caffeine evident on the 120min timepoint). It has previously been reported, that *ssp2* mutants display increased resistance to Torin1 because of increased TORC1 activity (Atkin et al., 2014). As *gsk3* genetically interacts with *ssp2* (Qingyun et al., 2016; Rallis et al., 2017), we investigated its role in mediating the activity of caffeine and Torin1 in relation to AMPK-related genes *ssp1, ssp2 and amk2*. Surprisingly, deletion of *gsk3* partially suppressed the cell cycle effects of Torin1 but not caffeine (Figure 5C). Deletion of both *gsk3* and *ssp2* strongly suppressed the effects of both caffeine and Torin1 on cell cycle progression (Figure 5D). Additionally, caffeine failed to advance cell cycle progression in *gsk3*Δ *ssp1*Δ and *gsk3*Δ *amk2*Δ double mutants (Figure 5E). We conclude that Ssp2 signalling mediates the effects of caffeine on cell cycle progression. In contrast, both Gsk3 and Ssp2 are required to mediate the effects of Torin1 on cell cycle progression. The intrinsic levels of TORC1 activity within *gsk3* and *ssp2* mutant cells have been presumed to be higher compared to *wt* cells (Rallis et al., 2017). Hence, both genes are involved in advancing mitosis in *S. pombe* but with differential effects on cell cycle dynamics. Mutants lacking Tor1 have been reported to be sensitive to caffeine (Rodríguez-López et al., 2023). We also noted that *tor1*Δ and *gad8*Δ mutants were sensitive to caffeine, suggesting this pathway confers resistance to the drug (Figure 5F, Supplementary figure 1F) Caffeine may thus impact on Gsk3 function via activation of TORC2 signalling albeit indirectly (Du et al., 2016). Indeed, TORC1 and TORC2 exert opposing roles on mitosis with the latter playing a role in cell cycle re-entry following DNA damage repair (Ikai et al., 2011).

**Figure 5.**
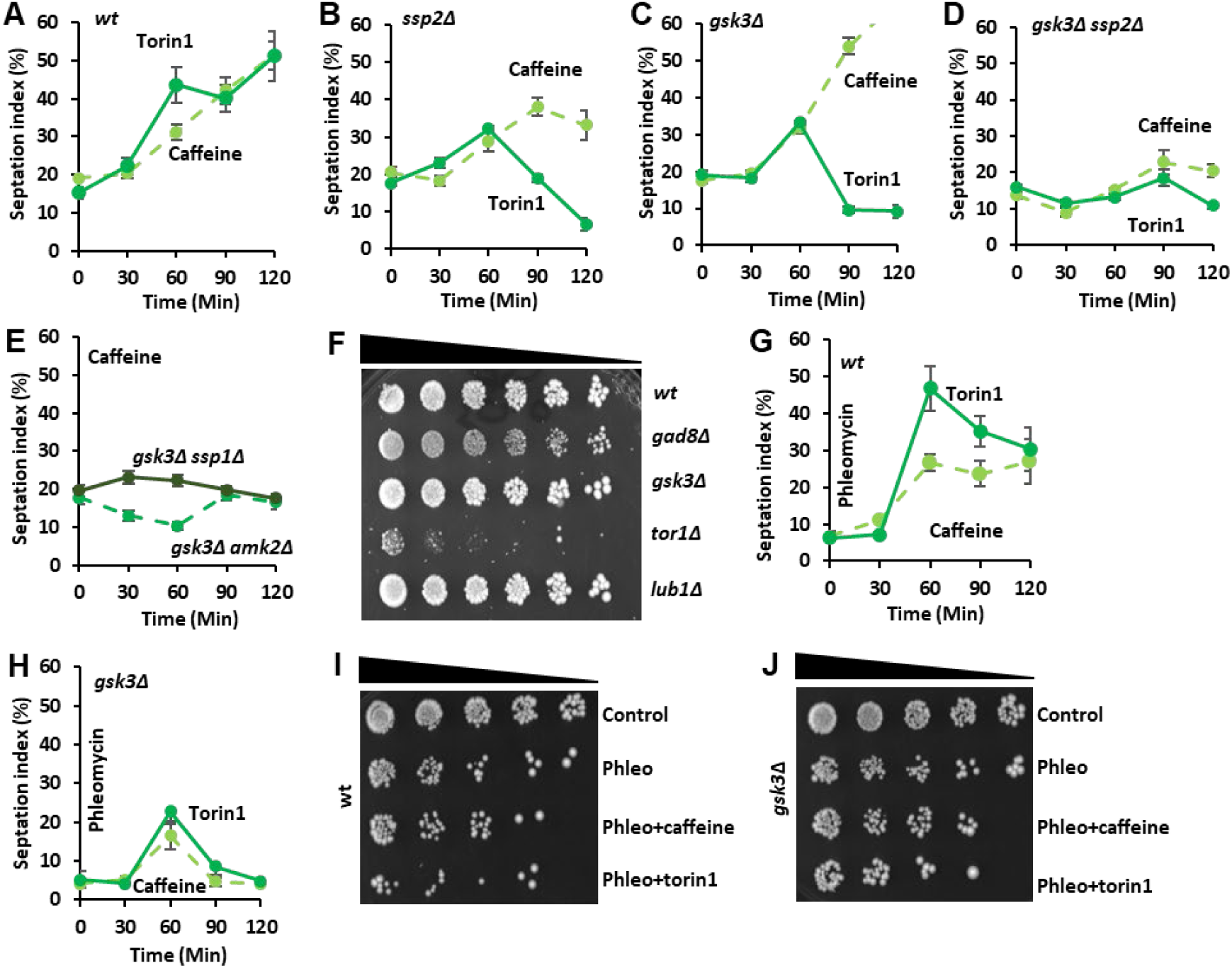
Caffeine modulates cell cycle kinetics via TORC1 and TORC2 complexes in *S. pombe*. **(A-E).** Cultures were incubated for 2h with or without 10mM caffeine or 5µM Torin1 in 10mL medium as indicated. Fixed cells were stained with calcofluor and the septation index determined using fluorescent microscopy. **(F).** The indicated wildtype and mutant strains were grown to log phase and resuspended in fresh YES media. Cultures were adjusted for cell number, serially diluted 3-fold and plated on YES agar containing 10mM caffeine. **(G-H).** Wildtype and *gsk3*Δ cultures were grown to log phase, left untreated or exposed to 5 µg/mL of phleomycin for 2h. Cultures were then exposed to 10mM caffeine or 5µM Torin1, harvested at the indicated time points and septation indices monitored. **(I-J).** Wild type and *gsk3*Δ cultures were grown to log phase, left untreated or exposed to 5µg/mL of phleomycin for 2h. Cultures were then exposed to 10mM caffeine or 5µM Torin1 for 2h, adjusted for cell number, serially diluted 3-fold and plated on YES agar.

We compared the effects of caffeine and Torin1 on cell cycle progression in *wt* and *gsk3*Δ mutants exposed to phleomycin. The effect of caffeine and Torin1 on DNA damage sensitivity was partially supressed in *gsk3*Δ mutants (Figure 5I and 5J, more evidently in the case of Torin1 in the shown conditions). It is unclear, why Gsk3 mediates the effects of caffeine in cells previously arrested with phleomycin but not under normal conditions. Gsk3 positively regulates the stability of the E3 ligase Pub1 which in turn regulates Cdc25 stability (Nefsky and Beach, 1996). Increased TORC1 activity in *gsk3*Δ mutants (Qingyun et al., 2016; Rallis et al., 2017) and strong inhibition of Cdc2 in the presence of phleomycin may attenuate the ability of caffeine to advance mitosis. These observations further demonstrate that differential mechanisms control the respective effects of caffeine and Torin1 on cell cycle progression via direct and indirect activation/ inhibition of TORC1 and 2.

### Caffeine exacerbates the *ssp1*Δ phenotype under environmental stress conditions

Mutants lacking *ssp1* respond weakly to the mitotic effects of caffeine (Figure 1B). Ssp1 has been reported to regulate cell cycle re-entry, following Sty1-mediated Cdc25 inhibition, by suppressing Srk1 expression (Gómez-Hierro et al., 2015). As Srk1 attenuates the effect of caffeine on cell cycle progression in a Sty1-dependent manner (Alao et al., 2014) we investigated its effect on the *ssp1*Δ mutant strain under heat and osmotic stress conditions. *ssp1*Δ mutant cells exposed to 1M KCl for 4h and 46h became elongated (cell size at division) from 14.3µm and 13.4µm to 16µm and 25µm at 4 and 46h respectively. Exposure to 10mM caffeine alone for 4h and 46 h also increased the average cell length from 14.3µm and 13.4µm to 18.1µm and 27.2µm respectively (Figure 6A). Microscopic analyses demonstrated, that *ssp1*Δ mutants are longer than *wt* cells following exposure to phleomycin (29.2µm versus 35.4µm) (data not shown). *ssp1*Δ mutants exhibited slight sensitivity when stationary phase cells were plated on YES media containing 10mM caffeine. In contrast, *amk2* Δ mutants were unable to regrow showing that caffeine exacerbated the phenotype of *ssp1*Δ mutants exposed to environmental stress (Figure 6B). At 35°C *ssp1*Δ mutants delay progression through mitosis (Gómez-Hierro et al., 2015). This effect was clearly exacerbated by co-exposure to 10mM caffeine (19.5µm versus 37.8µm after 24h). In contrast, both Rapamycin (100ng/mL) and Torin1 (5µM) were sufficient to overcome the *ssp1*Δ phenotype under these conditions (15.4µm and 21.8µm at 24h for Rapamycin and Torin1 respectively) (Figure 6C). Inhibition of TORC1 with Rapamycin or deletion of *sck1*, has been shown to permit growth under restrictive temperatures (Morozumi et al., 2024). The Ssp1-Ssp2-TORC1 pathway is required for proliferation in the presence of KCl. Caffeine enhanced the sensitivity *ssp1*Δ and *amk2*Δ mutants to KCl (Figure 6D). Caffeine at 10 mM and KCl at 0.6 M induced mitochondrial fission as seen through patterns of Cox4-GFP (Figure 6E), in a way like the reported effect of 1M KCl treatment (Schutt and Moseley, 2017). Together, these observations further suggest a role for the Ssp1-Ssp2-TORC1 pathway in mediating the effects of caffeine on cell cycle progression under normal and environmental stress conditions.

**Figure 6.**
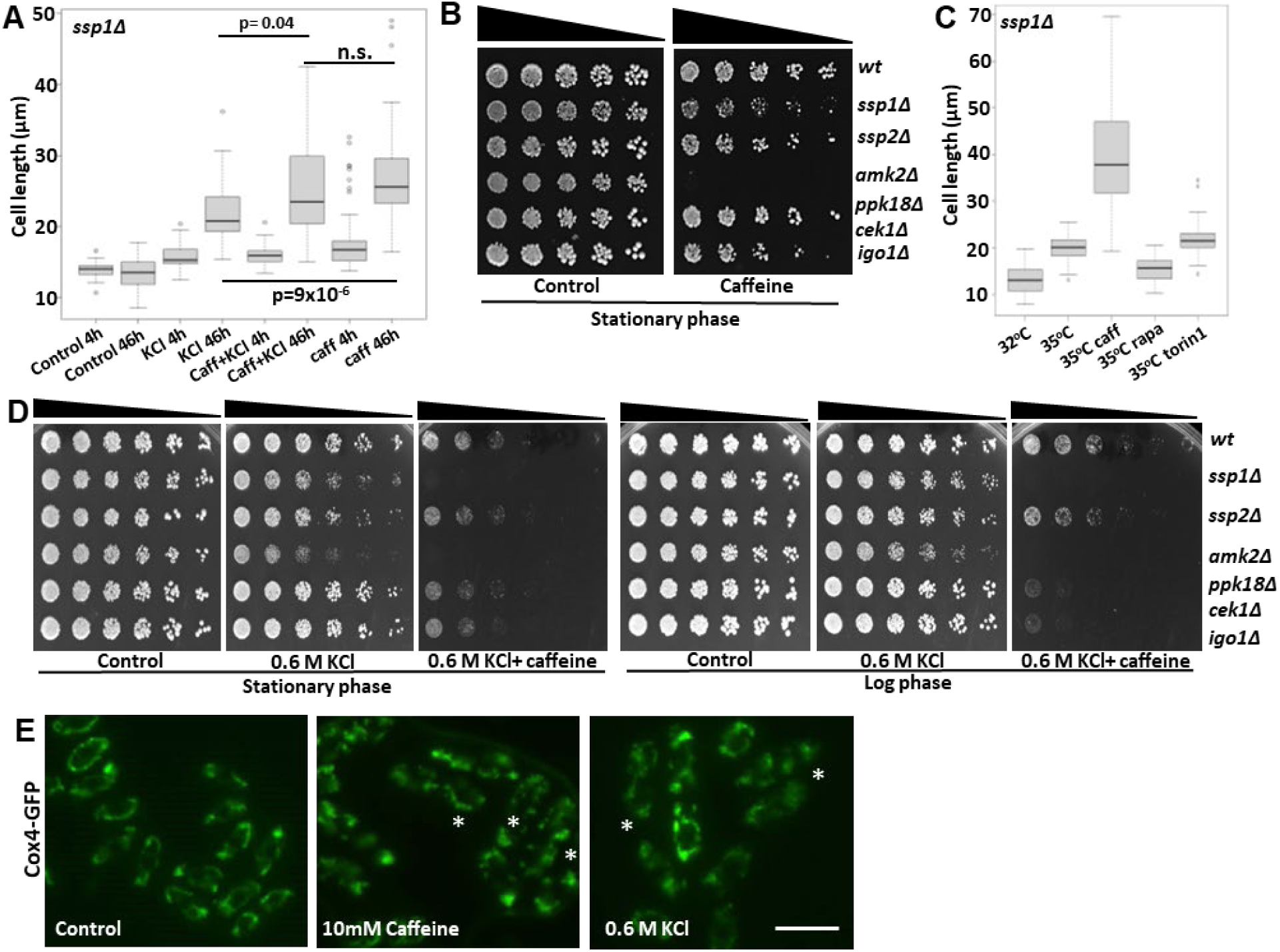
Caffeine signals via the Ssp1-dependent stress response pathway. **(A).** *ssp1*Δ mutants were grown to log phase at 32°C, resuspended in fresh YES media with or without 1M KCl or 10mM caffeine alone or in combination for 4h or 46 h. Cells were stained with calcofluor and the cell length determined using fluorescent microscopy (*n*=50). Under conditions where septate cells were absent, the average cell length of non-septate cells in at least 5 different fields was determined. **(B).** The indicated wildtype and mutant strains were grown to stationary phase, adjusted for cell number, serially diluted 3-fold and plated on YES agar containing 10mM caffeine. **(C).** *ssp1*Δ mutants were grown for 24h under the indicated conditions, harvested and treated as in A. **(D).** The indicated wild type and mutant strains were grown to stationary phase at 32°C. Cultures were adjusted for cell number, serially diluted 3-fold and plated on YES agar containing 0.6M KCl alone or in combination with 10mM caffeine. Alternatively, stationary phase cultures were adjusted to an OD_600_ of 0.3, grown to log phase and treated as in the stationary phase panel. **(E).** Cells expressing Cox4-GFP were exposed to 10mM caffeine or 0.6M KCl and mitochondria visualised by fluorescent microscopy. Asterisks indicate fragmentated mitochondria.

Amk2 has been reported to play a role in DNA damage repair and chromatin remodelling in *S. pombe* and mammalian cells (Ait-Saada et al., 2019; Lee et al., 2017). As caffeine is itself a genotoxin (Calvo et al., 2009; Rowley and Zhang, 1999), we further investigated the role of Amk2 in mediating resistance to caffeine. Amk2 specifically mediated the resistance of stationary phase cells plated on caffeine to the drug (Figure 6B and Supplementary figure 4A). When stationary phase cells were inoculated into YES media containing caffeine, *amk2* mutants showed a significantly reduced O.D. relative to *wt* cells over the first 24h. At 48h the *amk2*Δ and *wt* cultures had similar O.D.s suggesting *Amk2* regulates cell cycle re-entry or proliferation rate in the presence of caffeine (Supplementary figure 4B). This role is specific for Amk2, as we did not observe delayed proliferation in *ssp1*Δ, *ssp2*Δ, *cek1*Δ *ppk18*Δ and *igo1*Δ mutants. Given the specific sensitivity of *ssp2*Δ and *amk2*Δ mutants to phleomycin and their enhanced sensitivity to HU in the presence of caffeine (Figure 2J), Amk2 may mediate resistance to the DNA damaging effects of caffeine in stationary phase cells. When inoculated into media containing caffeine *wt* cells had approximately a 3-fold higher O.D. relative to amk2 mutants after 24h (Supplementary figure 4B and 4C).

### Caffeine enhances DNA damage sensitivity independently of its cell cycle effects

The E2 ubiquitin-conjugating enzyme Rhp6 (belonging to the histone H2B-K119 ubiquitin ligase complex HULC) cooperates with Cut8, a tethering factor of the nuclear proteasome, towards mediating DNA repair in *S. pombe*. Furthermore, Rhp6 has been shown to mediate the cell cycle effects of caffeine (Rowley and Zhang, 1999). Caffeine may thus interfere with Rhp6- and Cut8-mediated DNA damage repair (Kearsey et al., 2007; Rowley and Zhang, 1999; Takeda and Yanagida, 2005). We have confirmed that caffeine even at 10mM did not drive *rhp6*Δ mutants, previously exposed to HU, into mitosis (Figure 7A). In contrast, caffeine blocked the HU-induced cell cycle arrest in *cut8*Δ mutants (Figure 7B). While *rhp6*Δ mutants are checkpoint competent they were still sensitised to HU by caffeine (Figure 7C and 7E). We observed that *rhp6*Δ mutants became elongated indicating checkpoint arrest but remained sensitive to HU. Caffeine similarly enhanced the sensitivity of *cut8*Δ mutants to HU (Figure 7B and 7D). Caffeine may induce a specific form of DNA damage that is repaired by Rhp6 and enhances HU sensitivity (Calvo et al., 2009; Rowley and Zhang, 1999). Both *rhp6*Δ and *cut8*Δ mutants were highly sensitive to phleomycin with the latter losing viability rapidly (as noted by the lack of cell growth in the presence of phleomycin compared to *rhp6*Δ mutants) (Figure 7C to 7E). Caffeine alone or in combination with Rapamycin as well as Torin1 suppressed Cut8 expression in cells previously exposed to phleomycin but not under normal growth conditions (Figure 7F). These findings provide further evidence for a cell cycle-independent effect of caffeine on DNA damage sensitivity.

**Figure 7.**
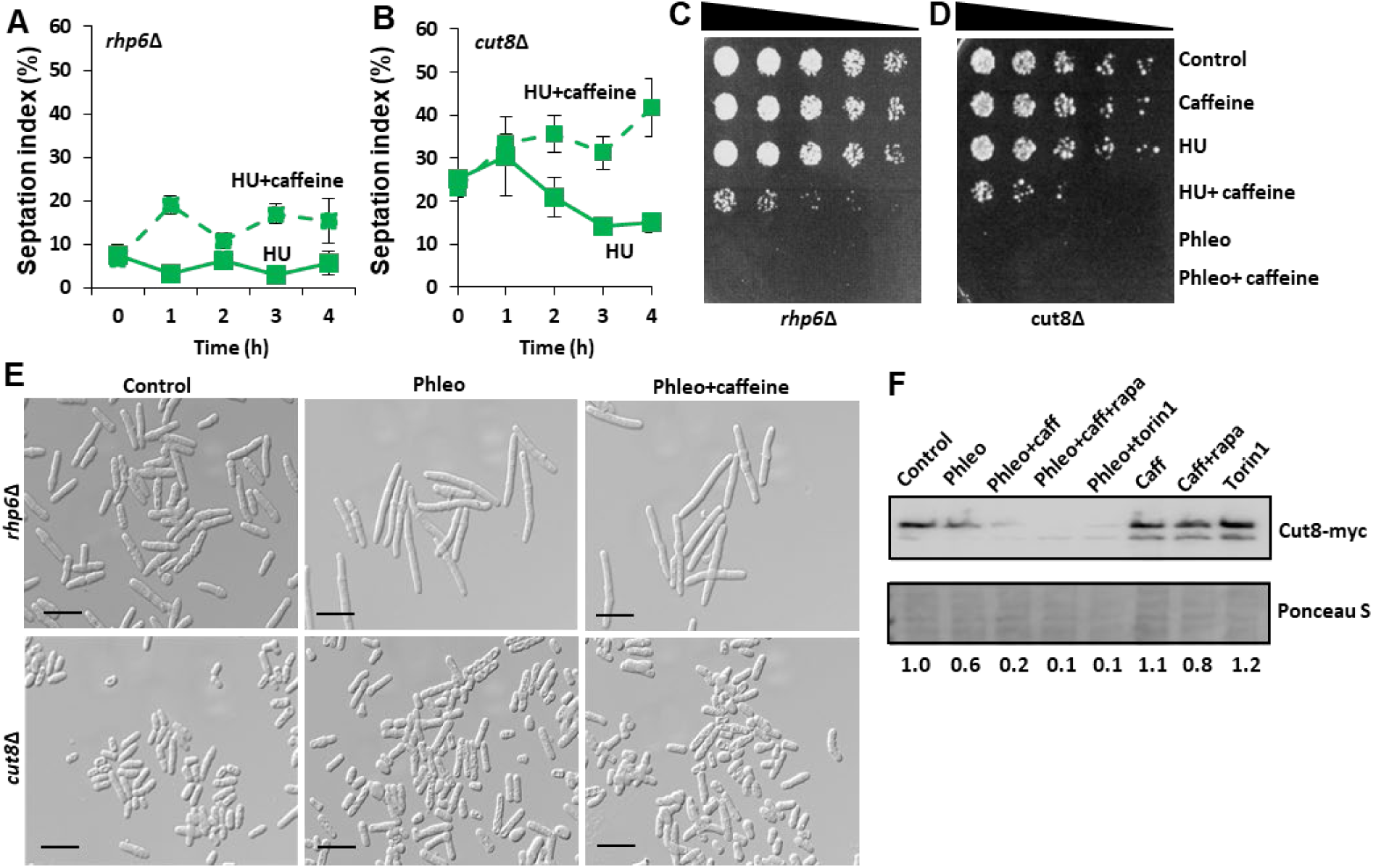
Caffeine enhances DNA damage sensitivity independently of checkpoint signalling and interferes with DNA damage repair. **(A and B).** *rhp6*Δ and *cut8*Δ strains were grown to log phase overnight. Cells were incubated with 20mM hydroxyurea (HU) with or without 10mM caffeine. Cells were stained with calcofluor and the septation index determined using fluorescent microscopy. (C and D). *rhp6*Δ and *cut8*Δ strains were grown to log phase and exposed to 20mM HU or 5µg/ mL phleomycin for 2h. Cells were then incubated for a further 2h following the addition of 10mM caffeine as indicated. Samples were adjusted for cell number, serially diluted and plated on YES media. **(E).** Samples of *rhp6*Δ and *cut8*Δ strains in C and D were examined by DIC microscopy. **(F).** Cells expressing Cut8-myc were left untreated or incubated with 5µg/ mL phleomycin for 2h. Cells were then incubated for a further 2h following the addition of 10mM caffeine, caffeine with 100ng/ mL Rapamycin or 5µM Torin1 as indicated. Samples were harvested, resolved by SDS-PAGE and Cut8 detected using antibodies directed against myc. Gel loading was monitored using ponceau S.

### The AMPK pathway is required for caffeine-mediated lifespan extension in *S. pombe*

We previously demonstrated that caffeine extends chronological lifespan (CLS) in *S. pombe* (Rallis et al., 2013). To determine if the Ssp1-Ssp2 pathway is required for caffeine-induced lifespan extension, we compared the effect of caffeine on CLS of *wt*, *ssp1Δ*, *ssp2Δ* and *amk2Δ* postmitotic (stationary phase) cell populations. Caffeine clearly extended CLS in *wt* cells (Figure 8A, log rank p=4.3×10^-4^). Caffeine failed to extend CLS in *ssp1Δ* (Figure 8B, log rank p=0.08), *amk2Δ* (Figure 9C, log rank p=0.85) and *ssp2Δ* (Figure 8D, log rank p=0.22). This result might be related with our earlier findings that these genes mediate resistance to caffeine (Figure 6). Although we cannot rule out, that caffeine is toxic to mutant cells under investigation, caffeine treatment is not detrimental to their chronological ageing (although a very mild, albeit not statistically significant effect, is observed for *ssp1*Δ).

**Figure 8.**
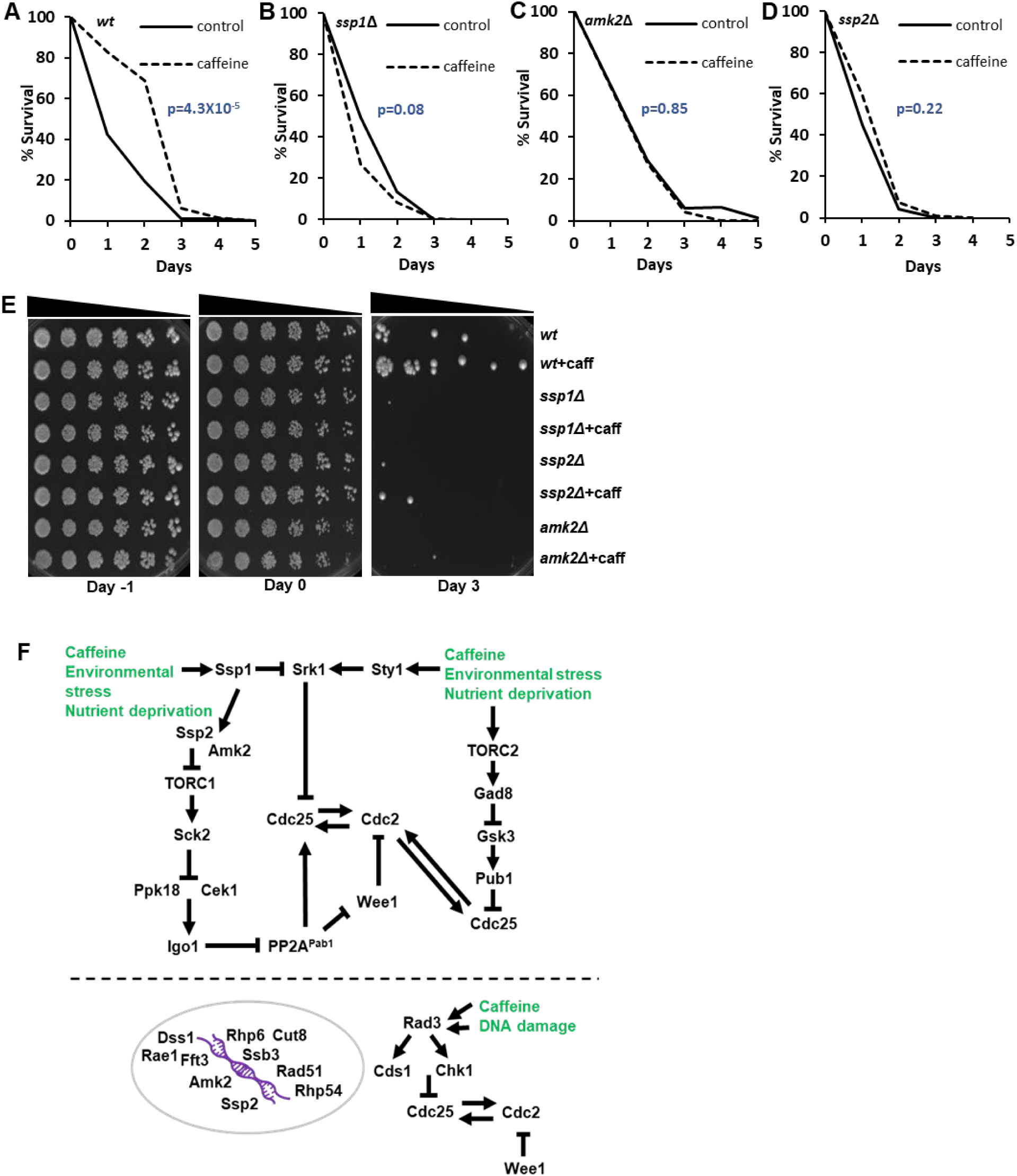
Caffeine extends lifespan in an Ssp1, Ssp2 and Amk2-dependent manner in fission yeast and summary of this work. **(A-D).** Wild type *ssp1*Δ, *ssp2*Δ and *amk2*Δ strains were grown to stationary phase with or without 10 mM caffeine as described in materials and methods. Statistical analyses represent the mean from at least 3 biological repeats (n= 3). **(E).** Semi-quantitative analysis of the effect of caffeine on CLS in fission yeast. The indicated strains were grown to log phase and then grown to stationary phase with or without 10 mM caffeine. Samples were harvested on the indicated days, adjusted for cell number (OD_600_ 0.3), serially diluted 3-fold, plated on YES agar. **(F).** Schematic model summarising the results of this study together with relationships from existing literature (see main text for details). Caffeine activates environmental stress response signalling and indirectly inhibits TORC1 advancing mitosis and modulating DNA damage repair signalling in fission yeast.

Semi-quantitative lifespan analyses using growing cells confirmed that *ssp1*Δ, *ssp2*Δ and *amk2*Δ mutants are short-lived relative to *wt* type cells. We performed these experiments to complement our standard CLS assays (Matecic et al., 2010). Cells were harvested at the indicated time points over a 4-day period to match our CLS assays, adjusted for cell numbers and plated on YES media (Figure 8E). Caffeine clearly extended the CLS of only *wt* cells consistent with a role for *ssp1*Δ, *ssp2*Δ and *amk2*Δ mutants in mediating resistance to caffeine (Figure 8A-8E). Together our findings suggest that all components of the *S. pombe* AMPK heterotrimer are required for caffeine-mediated extension of CLS.

## Discussion

In the present study, we sought to investigate if caffeine mediates its effects on cell cycle progression via the Ssp1 activated Ssp2-TORC1 signalling axis. In *S. pombe*, environmental stress and nutrient availability signalling converge via the TORC1 and TORC2 kinase complexes to regulate Cdc25 activity and cell cycle fate (Alao et al., 2023, 2021). We have previously demonstrated that caffeine drives mitosis by modulating Cdc25 and Wee1 activity in a manner reminiscent of TORC1 inhibition (Alao et al., 2020, 2014). Furthermore, caffeine modulates mitosis in *S. pombe*, by temporarily inhibiting Cdc25 activity via the Sty1-dependent activation of Srk1 (Alao et al., 2014; Calvo et al., 2009). However, caffeine stabilises Cdc25 independently of inhibitory phosphorylation sites (Alao et al., 2014) (Figure 4F and 4G).

Ssp1 is required for TORC1 inhibition and cell cycle re-entry following exposure to environmental stresses (Freitag et al., 2014; Gómez-Hierro et al., 2015; Schutt and Moseley, 2017). Our findings indicated that caffeine strongly interacts with Ssp1 to regulate mitosis and resistance to environmental stress. We also demonstrated that caffeine activates Ssp2 in an Ssp1-dependent manner. Ssp2 activation is required for the mitotic effects of caffeine in both log phase and DNA damage checkpoint-activated cell populations. Ssp2-mediated TORC1 inhibition relieves the inhibition of Sck2 on Ppk18 and Cek1, to drive cells into mitosis due to the downstream inhibition of PP2A^Pab1^ (Aono et al., 2019). Surprisingly *ppk18 cek1* co-deletion suppressed the mitotic effects of Torin1 but not those of caffeine. Thus, an additional kinase such as Ppk31 may mediate the effects of caffeine on mitotic progression. In contrast, *igo1* deletion suppressed the mitotic effects of both caffeine and Torin1. Igo1 is indispensable for rapid mitotic progression and proper cell cycle exit following TORC1 inhibition (Aono et al., 2019). Our findings suggest that caffeine mediates cell cycle progression in an Igo1-dependent manner, but this may be due to increased intrinsic PP2A^Pab1^ activity in *igo1*Δ mutants independently of Cek1/ Ppk18 (Figure 8F). It should be noted that caffeine differs from other stresses such as glucose deprivation and ionic stress by driving cells into mitosis, rather than delaying mitosis (Masuda et al., 2016), partially mimicking the effects of Rapamycin-induced TORC1 inhibition on cell cycle progression. However, signalling outputs downstream of TORC1 may differ between caffeine and direct TORC1 and/or TORC2 inhibitors. We noted for instance, that caffeine does not inhibit Maf1 phosphorylation but clearly suppresses Sck2 expression. 10mM caffeine has only a modest effect on cell growth in *S. pombe* compared to caffeine in combination with 100ng/mL Rapamycin or 5µM Torin1 exposure (Rallis et al., 2013; Rodríguez-López et al., 2020). Furthermore, ATP did not block the effects of caffeine on Ssp2 phosphorylation or mitosis. In contrast, ATP counteracted the negative effect of Torin1 on Ssp2 phosphorylation and mimicked the effect of *ssp1* or *ssp2* deletion on mitosis. In fact, addition of ATP mimicked the effect of ssp2 deletion on Torin1-induced mitosis. These observations suggest that caffeine activates Ssp2 independently of intracellular ATP levels. Caffeine induced mitochondrial fission in a manner similar to KCl as previously reported with the effect of KCl occurring independently of Ssp2 (Schutt and Moseley, 2017).

FACS analyses suggest that while caffeine drives cells into mitosis in an Ssp1-dependent manner, DNA replication may not be completely inhibited in *ssp1* mutants co-exposed to phleomycin. Deletion of *ssp1*, thus, may uncouple cytokinesis from DNA replication when these mutants are exposed to caffeine in the presence of phleomycin. Aside from *gsk3* mutants, we did not detect a differential decrease in DNA damage sensitivity in any of the Ssp1-Ssp2-TORC1 axis mutants. Interestingly, *ssp2*Δ and *amk2*Δ mutants showed enhanced sensitivity to phleomycin and were more sensitive to the effects of caffeine in the presence of HU. Caffeine-mediated activation of Ssp2 and Amk2 may thus have a protective effect during activation of the DNA damage response. Caffeine is clearly genotoxic in *S. pombe* at 10mM. Resistance to caffeine requires Rad3 and the Homologous Recombination (HR) DNA Damage Repair proteins Rad51 and Rhp54 as well as the 26S proteasome nuclear tether Cut8 required for DNA damage repair (Calvo et al., 2009; Rowley and Zhang, 1999). Furthermore, caffeine appears to induce Cds1 and Chk1 signalling and clearly enhances phleomycin sensitivity even in stationary phase cells (Alao et al., 2014) (Figure 2h). Early studies have, indeed, identified caffeine as a genotoxic agent (Lehmann, 1972; Prakash, 1981; Rowley and Zhang, 1999; Ts’o and Lu, 1964). Our studies suggest that caffeine may be inducing a specific type of DNA damage in HU-exposed cells. We have confirmed that the *S. cerevisiae* RAD6 homologue Rhp6 is required for the effects of caffeine on the HU-induced DNA damage checkpoint. Rhp6 is a ubiquitin conjugating enzyme required for post replication repair (PRR), while also mediating mitotic progression (Rowley and Zhang, 1999). Crucially, caffeine enhanced the sensitivity of *rhp6*Δ mutants to HU despite its inability to advance mitosis under these conditions. Rhp6 regulates the abundance of the 26S proteasome within the nucleus by regulating the stability of its nuclear tether Cut8 (Takeda and Yanagida, 2005). We also noted that caffeine suppresses Cut8 expression in the presence of phleomycin. Caffeine may, thus, interfere with ubiquitin-dependent degradation within the nucleus in *S. pombe*. These findings are consistent with our previous finding that caffeine advances mitosis by stabilising Cdc25 within the nucleus. Our observation, that this activity is more evident in HU-treated than in phleomycin-treated cells hints at cell cycle phase specific effects. When Cdc25 activity is high (High Cdc2 activity), caffeine is readily able to drive cells into mitosis in a manner partially dependent on Ssp2. When Cdc25 activity is low, however, (low Cdc2 activity), caffeine requires the Ssp1-dependent activation of Ssp2 to drive cells into mitosis via inactivation of Srk1 and PP2A^Pab1^ (Chica et al., 2016b; Gómez-Hierro et al., 2015; Schonbrun et al., 2013) (Figure 8F). Enhanced Cdc2 activity in cells exposed to caffeine may be sufficient to drive DNA replication in the absence of mitosis in *ssp1*Δ and *ssp2*Δ mutants. The ability of caffeine to enhance phleomycin sensitivity even in stationary phase cells, suggests it might interfere with DNA damage repair. Additionally, PP2A^Pab1^ and other phosphatases in this family regulate DNA damage repair (Kim et al., 2019; Ramos et al., 2019). Caffeine and other TORC1 inhibitors may, therefore, drive cells into mitosis, while inhibiting their ability to mediate DNA damage repair and TORC2 crosstalk. TORC2 has been reported to regulate stress and DNA damage resistance by modulating chromatin remodelling, gene expression particularly in S-phase and exit from DNA damage cell cycle checkpoints. Caffeine may, thus, also impact on DNA damage sensitivity due to its effects on TORC2 signalling (Schonbrun et al., 2009).

In this study, we identified a role for the AMPK β-subunit Amk2 in specifically mediating the resistance of stationary phase *S. pombe* cells when plated on YES media that contain caffeine. The AMPK β-subunit Amk2 is required for heterotrimer formation and its subcellular localization (Valbuena and Moreno, 2012). Ssp2 regulates the localisation of transcription factors and transcriptional repressors in fission yeast. When activated, Ssp2 also suppresses the expression of genes involved in ATP-dependent processes such as DNA replication, DNA repair, translation and the expression of genes required for cell cycle exit and sexual development (Valbuena and Moreno, 2012). Ssp2 is not strictly required for progression through mitosis, however, it may enhance it. These reports are consistent with our finding, that caffeine can induce mitosis in *ssp2* mutants exposed to phleomycin when Gsk3 is expressed. A strong genetic interaction has been reported for *amk2* and *gsk3*, suggesting the latter may suppress TORC1 in a TORC2-dependent manner (Rallis et al., 2017). The AMPK β and γ subunits are not required for the induction of mitosis following nitrogen stress. In contrast, all three subunits appear to be required for cell cycle resumption following exposure to osmotic stress through KCl. This suggests that Ssp2 differentially modulates mitosis in a context-dependent manner (Davie et al., 2015; Forte et al., 2019; Rallis et al., 2013; Schutt and Moseley, 2017; Valbuena and Moreno, 2012). Its major function may be to regulate transcription in response to a variety of environmental stresses (Saitoh et al., 2015; Valbuena and Moreno, 2012). In contrast to its orthologue in *S. cerevisiae* (SNF1), Ssp2 appears not to be required for the induction of autophagy in response to glucose deprivation in *S. pombe* (Pérez-Díaz et al., 2023). Rather, AMPK/Ssp2 migrates to the nucleus, where it phosphorylates the Scr1 transcriptional repressor to induce its nuclear export (Saitoh et al., 2015). This facilitates the increased expression of high affinity hexose transporters in response to glucose deprivation (Saitoh et al., 2015). We did not observe any effect of caffeine on Ght5 expression (data not shown, separate manuscript in preparation). Conversely, ATP did not affect caffeine-induced Ssp2 phosphorylation or cell cycle progression. Caffeine did induce mitochondrial fission similarly to KCl. Indirect caffeine-induced Ssp2 activation may sufficiently inhibit TORC1 activity, to drive mitosis via the inhibition of Sck2 and Igo1. This would lead to caffeine stabilised Cdc25 activation and its subsequent degradation, to permit activation of the Septation Initiation Network (SIN) and exit from or further rounds of cell cycle progression (Chica et al., 2016b; Esteban et al., 2008; García-Blanco et al., 2019).

We have previously demonstrated, that caffeine-induces Cdc25 nuclear accumulation independently of DNA damage checkpoint-induced phosphorylation (Alao et al., 2014). Furthermore, we demonstrated in this study, that TORC2 is required for caffeine resistance. TORC2 regulates protein stability by regulating Gsk3-mediated stabilization of the Pub1 E3-ligase, which in turn regulates Cdc25 stability. Torin1-mediated TORC2 inhibition, has been shown to induce Pub1 stabilisation in a Gsk3-dependent manner. Caffeine-induced TORC2 activation would thus induce Cdc25 stability by suppressing Pub1 expression (Nefsky and Beach, 1996; Wang et al., 2022). Indeed, a major difference between caffeine and Torin1, is that the latter induces Cdc25 degradation in a manner like nitrogen deprivation (Figure 4F and 4G) (Ducommun et al., 1990; Nefsky and Beach, 1996). The present study also demonstrated that Gsk3, mediates the mitotic effects of caffeine in a *ssp2* mutant background. As noted above, we previously demonstrated a strong genetic interaction between *gsk3* and *ssp2* (Rallis et al., 2017). Caffeine-induced Ssp2 activation, may thus lead to the activation of accumulated nuclear Cdc25 via Igo1-dependent PP2A^Pab1^ inhibition. This would explain the inability of caffeine to induce mitosis in *igo1*Δ mutants. In addition, PP2A^Pab1^ inhibition could result in Wee1 inactivation and degradation (Alao et al., 2020; Lucena et al., 2017; Qiu et al., 2018; Smith et al., 2007). Our previous studies demonstrated that caffeine influences the stability of both Cdc25 and Wee1.

Importantly, we noted that the cell cycle effects of caffeine are distinct from its ability to enhance DNA damage sensitivity. Several studies have suggested that caffeine is a genotoxic agent (Lehmann and Kirk-Bell, 1972; Prakash, 1981; Ts’o and Lu, 1964). Caffeine can clearly enhance DNA damage sensitivity independently of its effects on mitosis. This is clearly observable in *rhp6* mutants which are defective in DNA damage repair (Figure 7). It has been suggested, that caffeine may interfere with DNA damage repair (Rowley and Zhang, 1999). Alternatively, caffeine could synergistically induce DNA damage in the presence of other genotoxins. This could lead to the activation of DNA damage response pathways that are incompatible with HU for instance (Bakuradze et al., 2016). We also noted that caffeine suppresses Cut8 expression in the presence of phleomycin. This suggests caffeine interferes with DNA damage repair, as *cut8*Δ mutants are extremely sensitive to phleomycin. *cut8*Δ mutants are also sensitive to caffeine, providing further evidence for its genotoxic activity (Calvo et al., 2009). The enhanced sensitivity of *ssp2*Δ and *amk2*Δ mutants to phleomycin suggests a role for these genes in the DNA damage response. of *S. pombe* AMPK is, therefore, required for resistance to caffeine and this is illustrated by the inability of the drug to extend CLS in this organism.

## Conclusion

The precise mechanisms by which caffeine exerts its effects on cell cycle progression, DNA damage sensitivity and CLS in *S. pombe* have remained obscure for many decades (Alao and Sunnerhagen, 2020; Moser et al., 2000; Rallis et al., 2013). A clearer understanding of the molecular pharmacology of caffeine, can draw links between genes that are important in cellular physiology and pathology (Alao et al., 2014; Alao and Sunnerhagen, 2020; Osman and McCready, 1998). We have demonstrated that caffeine accelerates mitosis in a manner partially dependent on Ssp2-mediated TORC1 inhibition. Deletion of *ssp2* prevents caffeine-mediated DNA damage checkpoint override but not DNA replication. In contrast, deletion of *ssp1* which positively regulates Cdc25 activity blocked both effects of caffeine in the presence of phleomycin. This activity is dependent on Igo1, which lies downstream of TORC1 and Gsk3. The latter, functions downstream of TORC2 to regulate protein stability and TORC1 activity (Pérez-Hidalgo and Moreno, 2017; Wang et al., 2022). Caffeine actions depend on TORC1-TORC2 crosstalk to drive DNA replication and mitosis via Cdc25. We also demonstrate that Ssp1 and Amk2 mediate the ability of caffeine to extend CLS. Paradoxically, while Ssp2 appears to mediate the ability of caffeine to override DNA damage checkpoint signalling, it is also required for DNA damage resistance. In the presence of other genotoxins such as hydroxyurea, caffeine may act synergistically to induce DNA damage or interfere with its repair. At the same time, activation of Ssp2 may exert a protective effect masking the effects of caffeine on DNA damage sensitivity. Our findings suggest that caffeine enhances DNA damage sensitivity independently of its effects on mitosis. It would be interesting to further compare the effects of direct Ssp2 activators and caffeine on mitosis, DNA damage sensitivity and ageing in *S. pombe*.

## Materials and methods

### Strains, media, and reagents

Strains are listed in Supplementary Table 1. Cells were grown in yeast extract plus supplements medium (YES) (Formedium, Hunstanton, United Kingdom). Stock solutions of caffeine and hydroxyurea (HU) (Sigma Aldrich, Gillingham, United Kingdom) (100 and 200mM respectively) were prepared in water stored at −20°C. Phleomycin was purchased from Fisher scientific (Loughborough, United Kingdom) or Sigma Aldrich as a 20mg/mL solution and aliquots stored at −20°C. Torin1 (Tocris, Abingdon, United Kingdom) was dissolved in DMSO (3.3mM) and stored at −20°C. For treatment with potassium chloride (KCl), 0.6M KCl was prepared in YES media with or without 10mM caffeine. ATP was obtained from Alfa Aesar, Fisher Scientific, and dissolved in water (100mM). Aliquots were stored at −20°C. Experiments on the effect of caffeine on HU sensitivity, we carried out as previously reported (Khan et al., 2020).

### Microscopy

Calcofluor white (Sigma-Aldrich), 4,6-diamidino-2-phenylindole (DAPI) staining and septation index assays were carried out as previously described (Alao et al., 2014). Images were obtained using a LEICA DMRA2 microscope using a 100x objective using the appropriate filter set. Alternatively, studies were performed using an Evos M5000 imaging platform (Thermofisher Scientific, Dartford, United Kingdom).

### Immunoblotting

Antibodies directed against phospho-Ssp2 (#50081) were purchased from Cell Signalling Technologies (Leiden, The Netherlands). Antibodies directed against GFP (11814460001 Roche) were purchased from Sigma-Aldrich (Gillingham, United Kingdom). Antibodies directed against V5 (sc-81594), HA (F7) (sc-7392) and Myc (9E10) (sc-40) were from Santa Cruz Biotechnology (Heidelberg, Germany). Secondary antibodies directed against mouse (ab205719) and rabbit (ab205718) IgG were purchased from Abcam (Abcam, Cambridge, United Kingdom). Cell pellets were prepared for SDS-PAGE and treated as previously reported (Alao et al., 2014). Where deemed appropriate, images were quantified using Image J software.

### Flow cytometry

Cells were harvested and stored in 70% ethanol at −20° C until analysis. After rehydration with PBS cells were permeabilised with 0.1% Triton, 1%BSA in PBS and stained with1ug/ml FxCycle violet stain (ThermoFisher, catalogue number F10347). The samples were processed on a Biorad ZE5 Analyser using the violet laser to detect FxCycle violet stain. Flow rate was slow and linear range of fluorescent intensity was used to capture changes in DNA content. The data were subsequently analysed using FlowJo 10.7.2.

### Chronological Lifespan Assays

CLS was performed as previously described (Rallis et al., 2013; Roux et al., 2006). Stationary phase cells (∼16 h) in YES media, were diluted into fresh media at an OD_600_ 0.2 and grown to an OD_600_ 0.35-0.5. Cultures were treated with water or 10mM caffeine for a further 24h. Appropriate serial dilutions were plated on YES media daily for 7 days and Colony Forming Units (CFUs) were counted. Survival curves were statistically analysed with Kaplan-Meier survival plots and log rank tests from at least 3 independent experiments. Semi-quantitative CLS assays were also carried out as previously described (Matecic et al., 2010).

## Author’s contributions

Conceptualisation and design of study: J.P.A., C.R. Experiments and data analysis: J.P.A., J.K., D.S. and C.R. Manuscript writing: J.P.A., J.B., D.S. and C.R. Funding acquisition: C.R., J.B. All authors have read, revised and approved the final manuscript.

## Acknowledgements

We are grateful to the Bahler, Moreno, Moseley, Nakashima, Takeda, Young and Petersen laboratories and the Yeast Genetic Resource Centre (YGRC) Japan for mutant *S. pombe* strains. We are grateful to James Briscoe for help and access to flow cytometry facilities and expertise. C.R. acknowledges funding from the Royal Society (Research Award grant number RGS∖R1∖201348), BBSRC (grant numbers BB/V006916/1, BB/V006916/2) and MRC (grant number MR/W001462/1). D.S. was supported by the Francis Crick Institute which receives its core funding from Cancer Research UK (CC001051), the UK Medical Research Council (CC001051), and Wellcome Trust (CC001051). The authors declare that they have no competing interests.

