## Supplementary Figures 1-4 and Supplementary Table 1 for "Dissecting the cell cycle regulation, DNA damage sensitivity and lifespan effects of caffeine in fission yeast"

### Supplementary figure 1

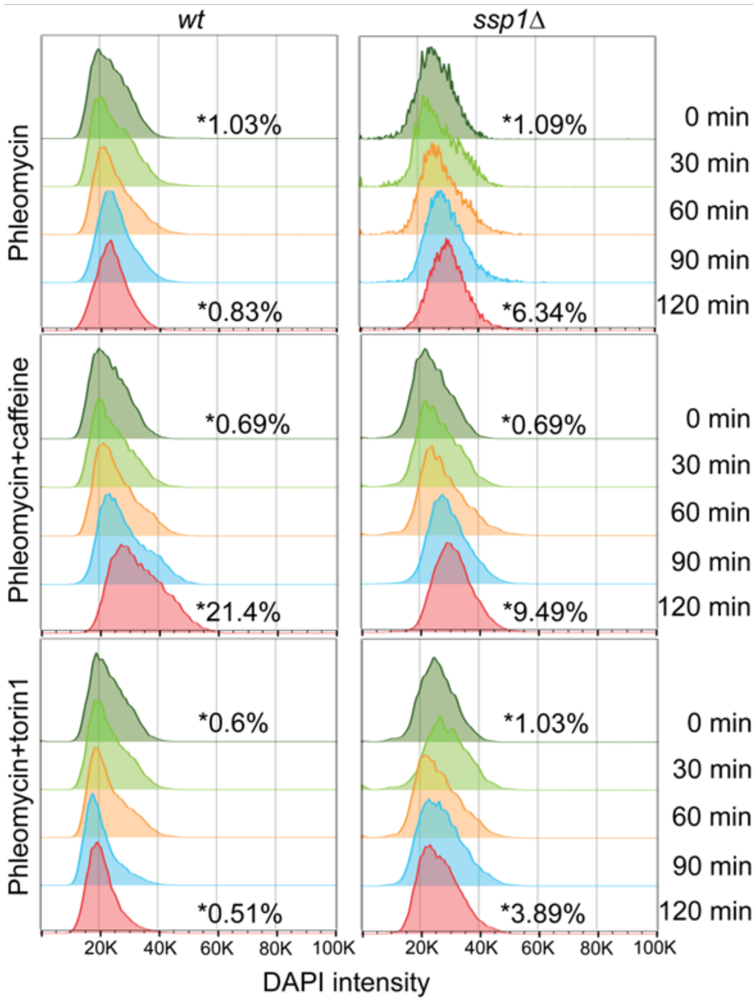


**Supplementary Figure 1. FACS analysis of *wt* and *ssp1* mutant cells following pharmacological treatments in a time course fashion as indicated.** *Wt* and *ssp1Δ* strains were exposed to 5μg/mL phleomycin for 2 hrs. Samples were then left untreated or co-exposed to 10mM caffeine or 5μM Torin1 and harvested every 30 minutes for 2 hrs. Numbers within the graphs indicate the percentage of cells that pass the 40K DAPI intensity arbitrary cutoff in each case.

number, serially diluted 3-fold, plated on YES agar and incubated for 3-5 days at 32°C.

**Supplementary figure 2**

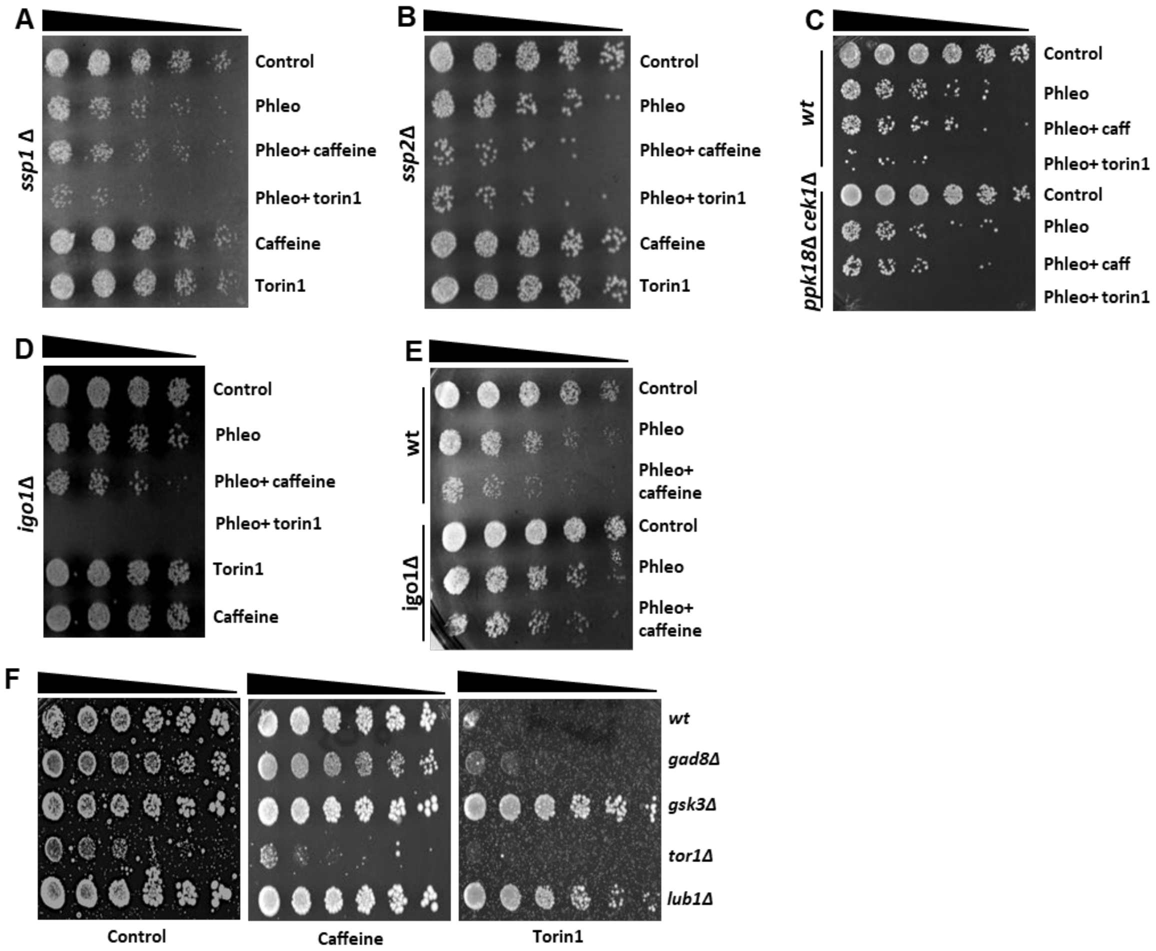


**Supplementary Figure 2. Caffeine enhances DNA damage sensitivity independently of its cell cycle effects. (A- E).** The indicated wild type and mutant strains were grown to log phase and left untreated or incubated with 5 µg/ mL phleomycin for 2h. Cultures were then incubated for a further 2h with or without 10mM caffeine or 5µM Torin1 in 10mL medium for a further 2h as indicated. Cultures were adjusted for cell. **F.** Tor1 mediates resistance to caffeine. The indicated wild type and mutant strains were grown to log phase and plated on media containing 10 mM caffeine or 5 µM Torin1.

**Supplementary figure 3**

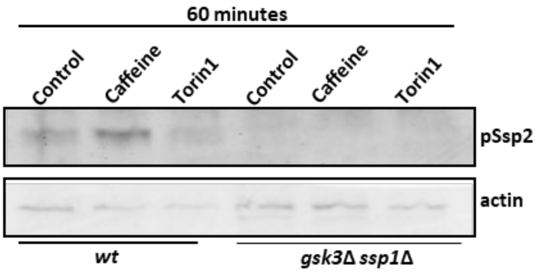


**Supplementary Figure 3. Validation of anti- phospho- Ssp2 antibody (Phospho- AMPKα (Thr172) (D4D6D) Rabbit mAb #50081)**. Wild type and *gsk3*Δ *ssp1*Δ mutants were incubated with 10 mM caffeine or 5 μM Torin1 for 60 minutes as indicated. Total cell lysates were resolved by SDS- PAGE and membranes probed with antibodies directed against phospho-Ssp2 and actin.

### Supplemental figure 4

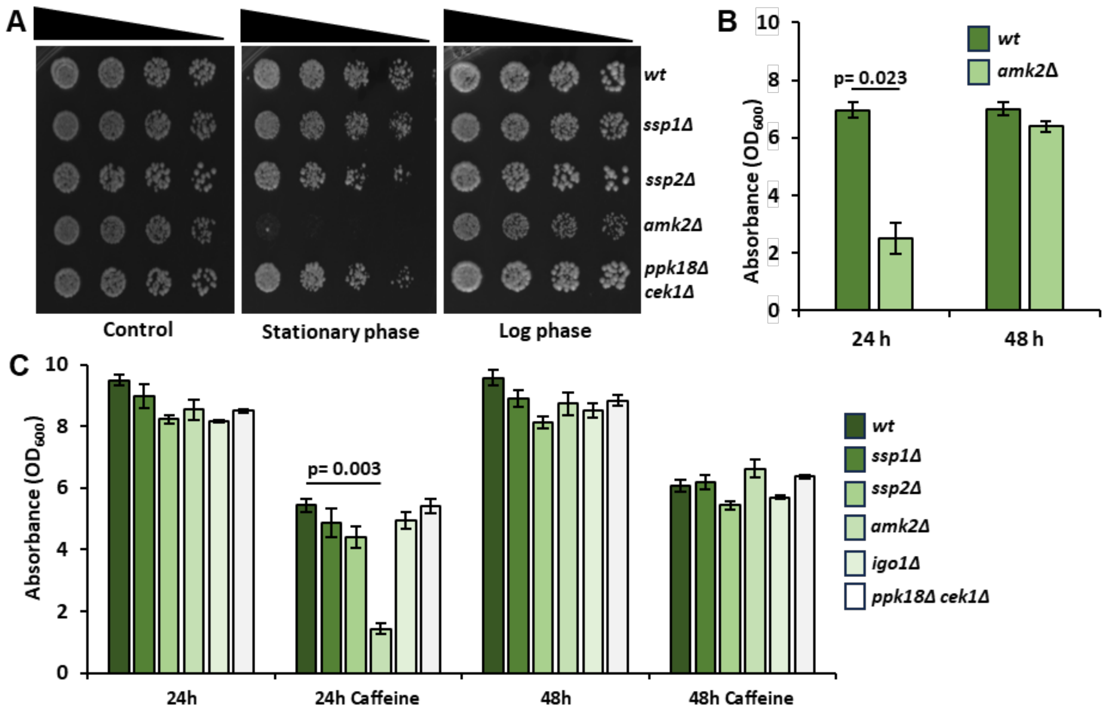


**Supplementary Figure 4. Amk2 is required for cell cycle re-entry in the presence of caffeine. (A).** The indicated wild type and mutant strains were grown to stationary phase. Cultures were adjusted for cell number, serially diluted 3-fold and plated on YES agar containing 10mM caffeine. Alternatively, cells were diluted into fresh media and grown to log phase before plating on 10mM caffeine. **(B).** Wild type and *amk2*Δ strains were grown to stationary phase and, diluted to an $OD_{600}$ of 0.2 and cultured in YES media containing 10mM caffeine for 24h or 48 h. Results represent the mean of at least 3 independent experiments. Error bars represent the mean ±S.E ($n= 3$). **(C).** Wild type and mutant strains were grown to stationary phase and treated as in B. Results represent the mean of at least 3 independent experiments. Error bars represent the mean ±S.E ($n= 3$).

**Supplementary Table 1.** List of fission yeast strains used in this study.

| Strain | Genotype | Source/Reference |
| --- | --- | --- |
| h⁻ L972 | h- | Laboratory stocks |
| tor1Δ | h- tor1::kanMX6 | Laboratory stocks/(Rallis et al., 2013) |
| gad8Δ | h- gad8:: ura4 ade6- M216 leu1 ura4-D18 | YGRC |
| ssp1Δ | h - ssp1::kanMX6 | Moseley laboratory/(Deng et al., 2017; Schutt and Moseley, 2017) |
| ssp1Δ | h - ssp1::kanMX6 | Laboratory stocks |
| pom1- GFP ssp1Δ | pom1-GFP-kanMX6 ssp1Δ::ura4 ura4-D18 | Bahler laboratory |
| ssp2Δ | h - ssp2::kanMX6, h + ssp2::kanMX6 | Moseley laboratory and Laboratory stocks/(Deng et al., 2017; Schutt and Moseley, 2017) |
| amk2Δ | h - amk2::kanMX6 | Moseley laboratory/(Deng et al., 2017; Schutt and Moseley, 2017) |
| gsk3Δ | h— gsk3::kanMX6 | Laboratory stocks/(Rallis et al., 2017) |
| gsk3Δ ssp1Δ | h— ssp1::kanMX6 gsk3::hphMX6 | Laboratory stocks/(Rallis et al., 2017) |
| gsk3Δ ssp2Δ | h— ssp2::kanMX6 gsk3::hphMX6 | Laboratory stocks/(Rallis et al., 2017) |
| gsk3Δ amk2Δ | h— amk2::natMX6 gsk3::hphMX6 | Laboratory stocks/(Rallis et al., 2017) |
| ppk18Δ | ppk18Δ::KanMX6 h+ | Moseley laboratory |
| cek1Δ ppk18Δ | h+ ppk18::KanMX6 cek1::ura4+ ura4-D18 | Moreno laboratory |
| cek1Δ ppk18Δ | h- Δcek1::natMX Δppk18::hphMX | Takeda laboratory |

| | | |
|---|---|---|
| igo1Δ | h- igo1Δ::KanMX6 | Moseley laboratory/(Deng et al., 2017; Schutt and Moseley, 2017) |
| rhp6Δ | | YGRC* |
| cut8Δ | h+ Δcut8::ura4+ | YGRC* |
| sck1-3HA | h+ sck1-3HA:natMX6 | YGRC* |
| sck1- 3HA | h-sck1+ -3HA: hphMX | Nakashima laboratory |
| sck2- 3HA | h-sck2+ -3HA: hphMX | Nakashima laboratory |
| ssp2- Myc | ssp2-13myc-hphR h+ | Moseley laboratory/(Deng et al., 2017; Schutt and Moseley, 2017) |
| cut8- 8Myc | h- leu1 cut8+- myc8 | YGRC |
| Maf1.pk | maf1.pk::KanMX6 | Petersen laboratory/(Davie et al., 2015) |
| cdc25- GFP | cdc25-GFPint cdc25:: kanMX6 ura4-D18 leu1-32 | Young laboratory |
| cdc25 (12A)- GFP | cdc25(12A)-GFPint cdc25:: kanMX6 ura4-D18 leu1-32 | Young laboratory |

*Yeast Genetic Resource Center (YGRC) Japan
